# Molybdate application in the early stages of shrimp growth suppresses sulphide formation in a shrimp pond bottom model

**DOI:** 10.1101/2024.09.25.614933

**Authors:** Funda Torun, Feyzâ Matisli, Barbara Hostins, Peter De Schryver, Nico Boon, Jo De Vrieze

## Abstract

Oxygen depletion and sulphide formation, resulting from the accumulation of organic waste, are common challenges in shrimp ponds that could result in complete harvest failure. The stage at which these circumstances occur during the shrimp growth period remains elusive, yet, knowledge of the timing of oxygen depletion and sulphide formation is essential to enable remediating actions. Here, we used an experimental shrimp pond model at different stages in the shrimp growth period to determine when oxygen depletion and sulphide production occur. Microscale depth measurements of oxygen and H_2_S were determined using microelectrodes to visualize their profiles at different depths of the water-sediment interface and the sediment. We evaluated the potential of different molybdate concentrations at different stages to determine the optimal conditions to suppress H_2_S formation. Oxygen depletion and sulphide production took place in the middle of the shrimp growth cycle in the simulated model of waste accumulation. The addition of molybdate was only effective in the early stages of the onset of oxygen depletion and H_2_S formation, and residual molybdate was required to ensure a continuous suppression of sulphide production. However, oxygen depletion could not be prevented and reintroduction of oxygen did not occur when molybdate was added. In conclusion, molybdate appeared to be an effective strategy to suppress H_2_S formation at the onset of its production in a shrimp pond bottom model.

## 1. Introduction

Shrimp farming is conventionally practiced in earthen outdoor ponds, where the post-larvae of shrimp reach their harvest size of about 15 g in a period of about 90 days (Hung and Quy, 2014). The pond bottom soil and the indigenously present and/or accumulated organics from earlier shrimp growth provide essential nutrients that contribute to the optimal conditions required to produce healthy and high-quality shrimp (Burford et al., 1998). Especially under semi-intensive and intensive culture conditions (50-300 shrimp m^-3^), a substantial amount of nutrients is coming from the feed applied, emphasizing the importance of its composition (Shiau, 1998). This pond bottom layer, thus, is an area that is densely populated by different groups of microorganisms, degrading the available organic matter into CO_2_ and other metabolites. Even though this is a natural process, which also occurs in natural water bodies, the oxygen consumption by aerobic microorganisms can cause a rapid drop in dissolved oxygen, both in the water column and sediment (Avnimelech and Ritvo, 2003). When oxygen consumption exceeds the oxygen supply, mainly due to intense microbial activity in the shrimp pond bottom from continued accumulation of organics through the shrimp feed, anaerobic conditions can arise (Dien et al., 2019).

The occurrence of anaerobic conditions, combined with the presence of sulphate, which is conventionally present in saline shrimp culture water, can give rise to the formation of hydrogen sulphide (H_2_S) from the activity of sulphate reducing bacteria (Avnimelech and Ritvo, 2003; Boyd, 1998; Kannan et al., 2018). These obligatory anaerobic sulphate reducing bacteria can use a plethora of organics or hydrogen as electron donor, with sulphate or S^0^ as electron acceptor, to produce H_2_S (Tang et al., 2009). In shrimp ponds, the formation of H_2_S can take place either in (1) so-called dead zones, *i.e.*, zones in the water column with insufficient mixing and, thus, insufficient oxygen supply from natural convection, or (2) in the pond bottom sediment. The formed H_2_S is toxic to the shrimp that dwell at the pond bottom, with both lethal effects, even at concentrations as low as 0.02 mg L^-1^ in Pacific white shrimp (*L. vannamei)* (US-EPA, 2011), and sublethal effects, such as lower diseases resistance and irritation of soft tissues, gut and stomach walls and deviant behaviour, at even lower H_2_S concentrations (Panakorn, 2016; Suo et al., 2017; Vismann, 1996). Hence, H_2_S formation is a critical cause of failed shrimp harvests, implying the urgent need for a targeted solution. However, the timing of applying such a solution is of critical importance, but at which stage of shrimp culturing oxygen depletion and H_2_S generation near the shrimp pond bottom occur, remains unclear.

Over the years, several strategies have been developed to deal with H_2_S toxicity in shrimp ponds. The sulphate reducing bacteria can grow over a broad range of pH (4.5-9.2) and temperature (3-70°C) values (Tang et al., 2009), so manipulating pH or temperature within the optimal values for shrimp aquaculture is impractical and unlikely to be successful. Nitrate addition could be an alternative strategy to control H_2_S formation in shrimp ponds. By adding nitrate, denitrifying bacteria can outcompete sulphate reducing bacteria in the competition for available carbon sources, thus, reducing H_2_S formation (Schwermer et al., 2010; Torun et al., 2020). However, adding nitrate might directly, through the formation of toxic metabolites, *e.g.*, nitrite (Ciji and Akhtar, 2020), or indirectly, by causing harmful cyanobacteria or algae blooms (Alonso-Rodrı guez and Páez-Osuna, 2003), induce toxicity to the shrimp. The H_2_S suppression mechanism of nitrate is also only temporary, because when nitrate is depleted, sulphate reduction and, thus, H_2_S production can recover (Schwermer et al., 2010; Torun et al., 2020).

A targeted strategy towards direct inhibition of sulphate reducing bacteria resides in the application of molybdate (MoO_4_), which is a sulphate analogue that can inhibit the microbial sulphate reduction metabolism that leads to the production of H_2_S (Nemati et al., 2001; Predicala et al., 2008). Recently, we discovered that molybdate could be used successfully to suppress H_2_S formation in shrimp pond models, both at the short-term (Torun et al., 2022) and throughout the entire shrimp culturing (Torun et al., 2024). However, since the sulphate reduction process depends on the availability of organic matter for the sulphate reducing bacteria, the relation between the load of organic matter and required molybdate concentration and timing of addition in the shrimp culturing that effectively prevents H_2_S formation in a shrimp pond bottom still poses a critical knowledge gap towards successful suppression of H_2_S formation in shrimp ponds.

In this study, we used an experimental set-up that simulates a shrimp pond bottom to investigate two key objectives. The first objective was to determine the stage in the shrimp culturing when oxygen depletion occurs and H_2_S production takes place in the pond water. The second objective was to unravel the suppressive effect of molybdate towards sulphate reduction, in function of different loads of organic waste accumulation, matching different stages in the shrimp culturing. The key hypothesis is that molybdate can be used as a targeted approach to mitigate sulphide production in shrimp pond bottoms at the appropriate moment during shrimp culturing.

## 2. Material and methods

### 2.1. Sampling and storage

#### 2.1.1. Sediment sampling and preparation

Sediment used in the different experiments was collected from the Ijzermonding Nature Reserve (Nieuwpoort, Belgium) from a creek (51°8′45′′ N/2°44′38′′E) that was regularly water-logged with tidal movement. Sediment was collected from the creek using a shovel after which the samples were transported to the laboratory in a closed plastic vessel. Prior to the experiments, the sediment was pre-treated by drying and ventilating the sediment at 28°C for a week, as described by Torun et. al. (2022). To avoid damage to the sensitive microprofiling equipment, and to remove large burrowing microfauna, the pre-treated sediment was also sieved over 1 mm, and stored at 4°C until use. The sediment was prepared for the experiments using a standard procedure for aquaculture pond sediments (Thunjai et al., 2001). Briefly, the sediment was dried at 105°C, pulverized and sieved over a 1-mm sieve after which it was diluted with distilled water in a 1:2 sediment:distilled water ratio.

#### 2.1.2. Feed and faeces collection and storage

Shrimp faeces were collected directly from the flush outlet of shrimp tanks where *Litopenaeus vannamei* at post-larvae phase were fed with CreveTec Grower 2 (CreveTec, Ternat, Belgium), at the facilities of the Aquaculture and *Artemia* Reference Center, Ghent University, Belgium. For each experiment cycle, a fresh batch of faeces was collected to avoid organic matter degradation during storage. The pH, conductivity and total solids (TS) content of the faeces were measured directly upon arrival in the laboratory. The CreveTec Grower 2 was used as dosed feed, and was stored at −20 °C until addition.

### 2.2. Experimental set-up and operation

An experimental set-up that allowed mimicking the conditions in the pond sediment in shrimp ponds as closely as possible was established (Figure S1). To ensure controlled conditions, no shrimp were included in the set-up to maintain the focus on the microbial processes associated with H_2_S formation. All experiments were carried out using this simplified pond bottom model. Prepared sediment (section 2.1.1) was added to 250 mL size glass beakers (outer diameter 70 mm), to a sediment depth of 3.5 cm (total 100 g of prepared sediment for each beaker) to simulate an earthen pond bottom. The sediment layer was overlaid with 5 cm artificial seawater (Instant Ocean, Aquarium Systems, Mentor, OH, USA). The salinity of the artificial seawater was adjusted to 20 g L^-1^ to represent a common salinity in shrimp ponds (15-25 g L^-1^). To avoid excessive water evaporation, all beakers were put in a transparent large plastic box with a non-airtight lid that was kept closed until microscale depth profiles or water column measurements were performed. This strategy was applied to reduce water evaporation to a negligible extent, while ensuring the availability of oxygen. A light regime of 12 h light on/off using a fluorescent lamp was applied, and no active aeration was provided. To balance the gradients at the sediment-water interphase, the sediment and overlying water were kept in a temperature-controlled room at 28 ± 1°C for 24 hours after adding the sediment and overlaying water, before adding the waste (feed and faeces) the next day. To determine the amount of waste added, semi-intensive stocking of 50 shrimp m^-2^ was used as a reference. About 25% of cumulative input feed was assumed to be accumulating in the pond bottom, with 15% considered digested feed (faeces), while 10% was uneaten feed (Table S1). The day when waste was added was considered day 0 of the experiment. All experiments were carried out in the same temperature controlled room at 28 ± 1°C. Two sets of experiments were performed in this study.

#### 2.2.1. Experiment 1: Determination of the onset of oxygen depletion and H_2_S production

In the first experiment, by using the established shrimp pond model, the aim was to determine at which stage in the shrimp culturing oxygen depletion and H_2_S production occur in function of the accumulated waste loads. The 90-days period of culturing was divided into periods of 15 days (Table S2), resulting into seven different treatments, which were all carried out in biological triplicates, each reflecting a certain stage in the shrimp culturing, expressed in days of culture (DOC): DOC0 (control), DOC15, DOC30, DOC45, DOC60, DOC75 and DOC90. In each treatment, the total amount of waste accumulated over the previous culturing period was added in the form of feed and faeces, as illustrated in Table S1 & S2. The microscale depth profiles of O_2_, H_2_S and pH at the water-sediment interface and throughout the sediment were measured on day 0, 2, 4, 7 and 10.

#### 2.2.2. Experiment 2: Molybdate treatment at different stages of shrimp culturing

In the second experiment, the objective was to understand how the addition of molybdate would affect the H_2_S production in response to different organic waste loads, related to the stage in the shrimp culturing, *i.e.*, the DOC. Four different treatments were conducted, *i.e.*, DOC90 and 50 mg L^-1^ Na_2_MoO_4_.2H_2_O; DOC90 with 25 mg L^-1^ Na_2_MoO_4_.2H_2_O; DOC60 with 50 mg L^-1^ Na_2_MoO_4_.2H_2_O; and DOC45 with 25 mg L^-1^ Na_2_MoO_4_.2H_2_O treatment. The concentrations of Na_2_MoO_4_.2H_2_O were selected based on earlier research (Torun et al., 2022), and the DOCs were selected based on experiment 1.

For the addition of sodium molybdate at the appropriate concentrations, a 1 g L^-1^ Na_2_Mo_4_.2H_2_O stock solution was prepared by dissolving sodium molybdate (Sigma Aldrich, St. Louis, Mo., US) in an instant ocean solution at 20 g L^-1^ salinity. All experiments were run together with their respective control, *i.e.*, no molybdate addition, and in 6 biological replicates for 10 days. The O_2_ and H_2_S concentrations and pH were measured in the water column, about 1 cm above the water-sediment interphase, on day 0, 2, 4 and 7 and 10. These bulk measurements, instead of sediment profiles, of the O_2_ and H_2_S were selected, since H_2_S in the water column is the major concern for the shrimp that dwell on the pond bottom. After each microelectrode measurement (excluding day 0), one replicate from the control and one replicate from the treatment incubation were sacrificed to sample the water column to measure molybdate and sulphate concentrations. This sacrificial sampling was carried out to avoid the disturbance in water-sediment interface during the liquid sampling. The liquid samples were filtered over a 0.20 μm Chromafil^®^ Xtra filter (Macherey-Nagel, PA, USA), and stored in the fridge (4 °C), before the analysis of sulphate and molybdate concentrations.

### 2.3. Microelectrode measurements

The microscale depth profiles of O_2_, H_2_S and pH were measured with commercial microelectrodes (Unisense A.S. Denmark, tip sizes pH: 200 μm, H_2_S: 100 μm, and O_2_: 50 μm), operated with a motorized micromanipulator (Unisense A. S., Denmark). The oxygen profiles were measured separately at 500 μm resolution, while pH and H_2_S were simultaneously recorded with a 200 μm resolution at the water-sediment interphase, and also through the deeper layers of the sediments. The sensors were calibrated following a standard calibration procedure (Malkin et al., 2014). The O_2_ sensor was calibrated with a 2 point standard curve, using 100% in air bubbled seawater for the saturated dissolved oxygen concentration at 28°C and argon bubbled seawater for a dissolved oxygen concentration at zero. The H_2_S sensor was calibrated with a 3-5 point standard curve using a Na_2_S solution. The pH sensor was calibrated with 2 point calibrations using commercial (Carl Roth GmbH & Co.KG, Karlsruhe, Germany) pH buffer solutions at pH 4 and 7.

For the molybdate treatment experiments (experiment 2), the microelectrodes were utilized manually to take the O_2_, H_2_S and pH measurements at approximately 1 cm above the sediment surface, after ensuring that there was negligible variability in the duplicate measurements of the water column parameters.

### 2.4. Analytical techniques

The TS was determined according to Standard Methods (Greenberg et al., 1992). The pH of the overlaying water and sediment samples was measured with a standard pH meter (Methroohm, Herisau, Switzerland), which was calibrated using pH buffer solutions at pH 4 and 7. The conductivity was measured with a conductivity electrode (Consort C532, Turnhout, Belgium), which was calibrated with standard solutions of 0.01, 0.1 and 1 M KCl. The concentrations of sulphate were determined with ion chromatography (930 Compact IC Flex, Metrohm, Herisau, Switzerland), equipped with a Metrosep A supp 5–150/4.0 column with conductivity detector, after diluting the samples 1:50 using ultra-pure water (Milli-Q, Millipore Corparation, Burlington, MA, USA). The detection limit was 0.05 to 200 mg SO_4_ ^2-^ L^-1^. Molybdate concentrations were measured with a commercial kit (Hach, Model Mo-2, USA), which was based on the colorimetric determination of molybdenum using mercaptoacetic acid (Will and Yoe, 1953), specific for Mo^6+^ measurements. Standard solutions of 0, 5, 10, 25, and 50 mg L^-1^ Na_2_MoO_4_.2H_2_O were prepared to determine the calibration curve at 425 nm using a UV-Vis Spectrophotometer (WPA Lightwave II, Thermofisher, USA).

## 3. Results

### 3.1. Timing of oxygen depletion and H_2_S production during shrimp culturing

To determine at which stage in the shrimp culturing oxygen depletion and H_2_S production would take place, dissolved oxygen and H_2_S were monitored for 7 days under different waste load simulations, reflecting different stages in the shrimp culturing. In the water column, oxygen remained present throughout the 7 days of the experiment in the control treatment (DOC0) and DOC15 and DOC30 treatments (Figure 1). In DOC0, the average dissolved oxygen concentration in the water column dropped from 196 ± 2 µM on day 0 to 120 ± 5 µM on day 7, while it dropped from 193 ± 1 µM to 114 ± 2 µM in DOC15 and from 186 ± 3 µM to 119 ± 3 µM in DOC30, confirming that oxygen remained abundantly present. In contrast, in DOC45 the average dissolved oxygen concentration in the water column dropped from a value of 180 ± 2 µM on day 0 to below detection limit on day 1, and a similar observation could be made for DOC60, DOC75 and DOC90. Some (passive) reaeration occurred in DOC45 on day 5 and 7 to values of 77 ± 11 µM and 63 ± 6 µM, which was not the case in DOC60, DOC75 and DOC90. In the sediment, in the DOC0, 15 and 30 treatments, oxygen was detected in the first 3-4 mm of the sediment top layer after day 0. In the DOC45 treatment, despite partial reaeration, no oxygen could be observed in the sediment below 500 µm depth, and no oxygen was detected at all in the sediment in the DOC60, DOC75 and DOC90 treatments, which matched with the water column results.

**Figure 1.**
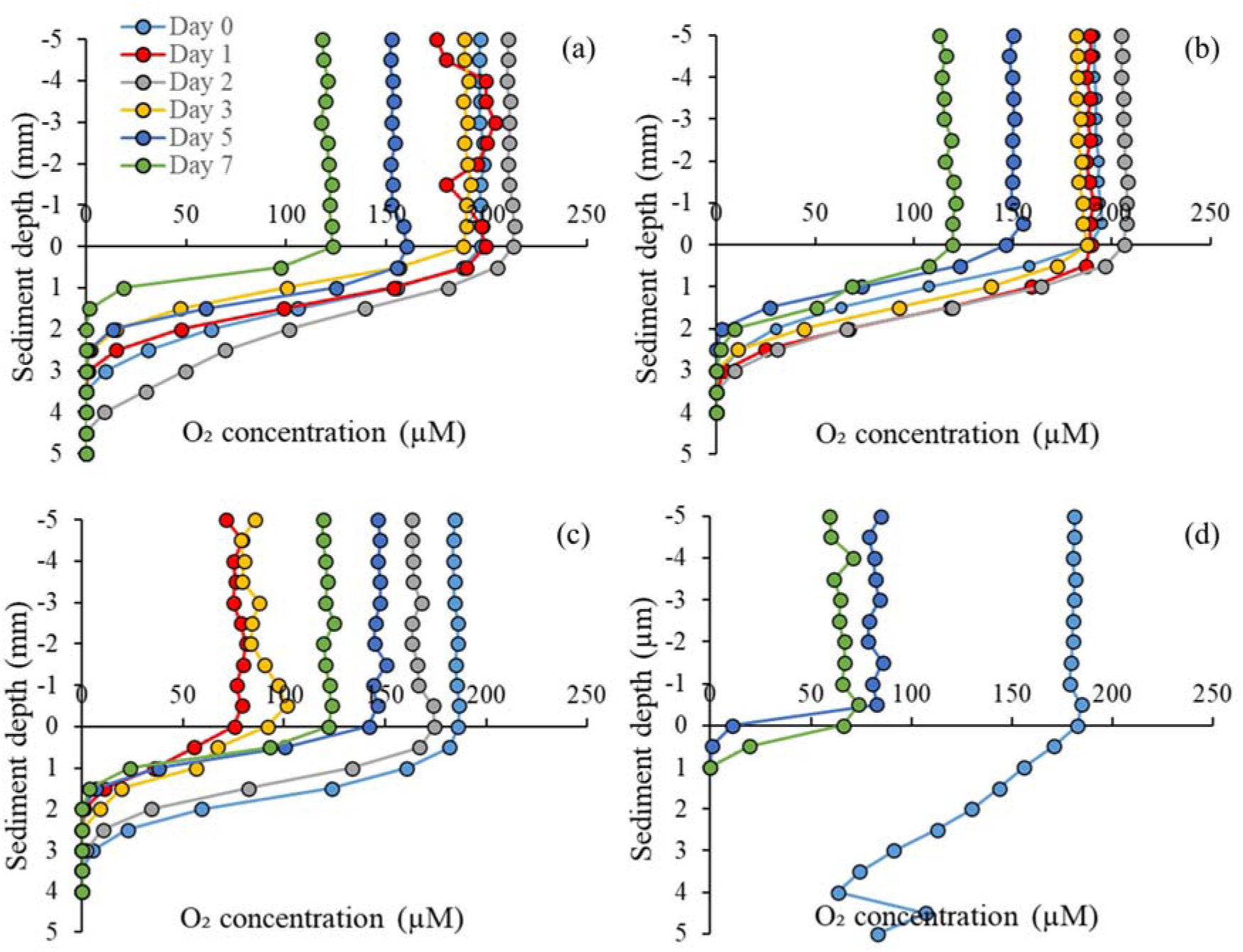
The O_2_ depth profiles for the (a) control or DOC0 (day of culture) treatment, (b) DOC15, (c) DOC30, and (d) DOC45. Values represent averages of biological triplicates, and error bars are omitted to maintain the visibility of the graphs. Zero depth equals to the sediment-water interface. No data for DOC60, DOC75 and DOC90 are presented, since, with the exception of day 0, no O_2_ was detected.

For the DOC0, 15 and 30, the H_2_S in the water column remained below 2 µM, related to the presence of oxygen, throughout the entire 7-day experimental period. In the DOC45 treatment, the H_2_S concentration increased to 3.0 ± 0.2 µM in the water column in the absence of oxygen on day 3 (Figure 2). In the DOC60 treatment, the H_2_S concentration in the water column increased to 11.8 ± 0.3 µM on day 3, while an even stronger increase up to 51 ± 4 µM H_2_S in DOC75, and 128 ± 2 µM in DOC90 was observed, both also on day 3. Like in the water column, the H_2_S concentration in the sediment remained below 2 µM in the DOC0, DOC15 and DOC30 treatments. In the DOC45 treatment, a H_2_S peak up to 11.9 ± 10.5 µM was measured on day 2 at a sediment depth of 1.6 mm, but on all other days, the H_2_S concentration remained below 5 µM. In the DOC60 treatment, the H_2_S concentration strongly increased in the upper 1 mm of the sediment layer to an average value of 11.4 ± 0.4 µM on day 3, which relates with the strong increase in H_2_S in the water column in DOC60 compared to DOC45. In the DOC75 and DOC90 treatments, this H_2_S peak in the sediment was much more substantial than in the DOC45 and DOC60 treatments, reaching a maximum value of 169 ± 95 µM at 2.4 mm depth in DOC75 and 389 ± 164 µM at 2.2 mm depth in DOC90, both on day 3.

**Figure 2.**
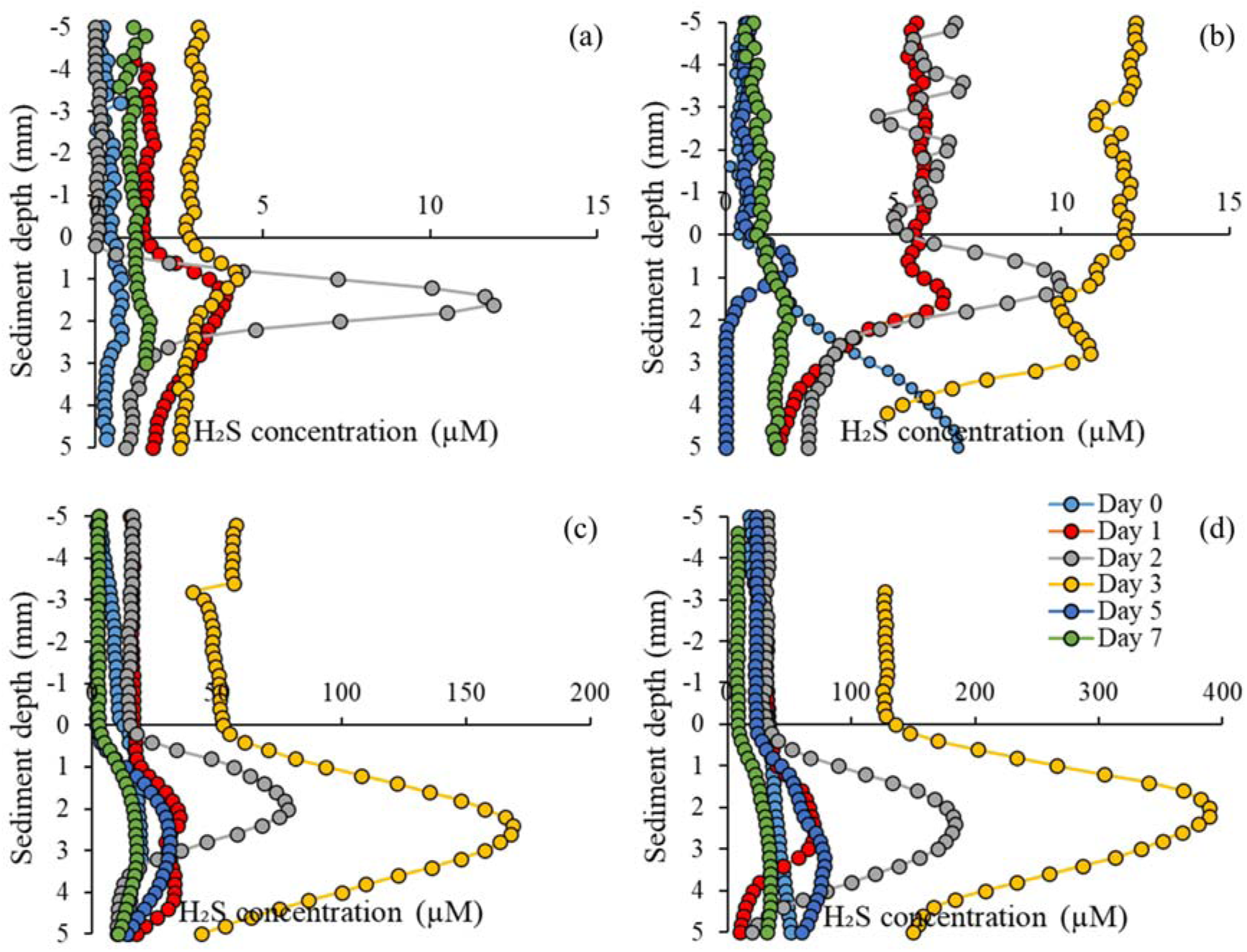
The H_2_S depth profiles for the (a) DOC45, (b) DOC60, (c) DOC75, and (d) DOC90 (day of culture). Values represent averages of biological triplicates, and error bars are omitted to maintain the visibility of the graphs. Zero depth equals to the sediment-water interface. No data for DOC0, DOC15 and DOC30 are presented, since H_2_S concentration remained below 2 µM, both in the liquid and bulk. Because of technical problems with the microelectrode, data from day 5 are not included for DOC45.

The pH in the water column showed two clear trends (Figure 3). First, the average pH in the water column on day 0 decreased with increasing waste load from 7.91 ± 0.00 in DOC0 to 6.30 ± 0.04 in DOC90. Second, while in DOC0 the pH decreased from 7.91 ± 0.00 on day 0 to 7.64 ± 0.01 on day 7, this decreasing trend was inverted in DOC30, with an increase in pH from 7.45 ± 0.01 on day 0 to 7.88 ± 0.00 on day 7. In the DOC90 treatment, this increase in pH over time was most substantial, raising from 6.30 ± 0.04 on day 0 to 8.03 ± 0.00 on day 7. In the sediment, in the DOC0, DOC15 and DOC30 treatments, pH decreased in function of sediment depth from day 1 on. In DOC45, the pH first increased and then decreased in function of sediment depth on day 1, and decreased in function of depth on day 2, 3, 5 and 7. A similar observation could be made for the DOC60, DOC75 and DOC90 treatments.

**Figure 3.**
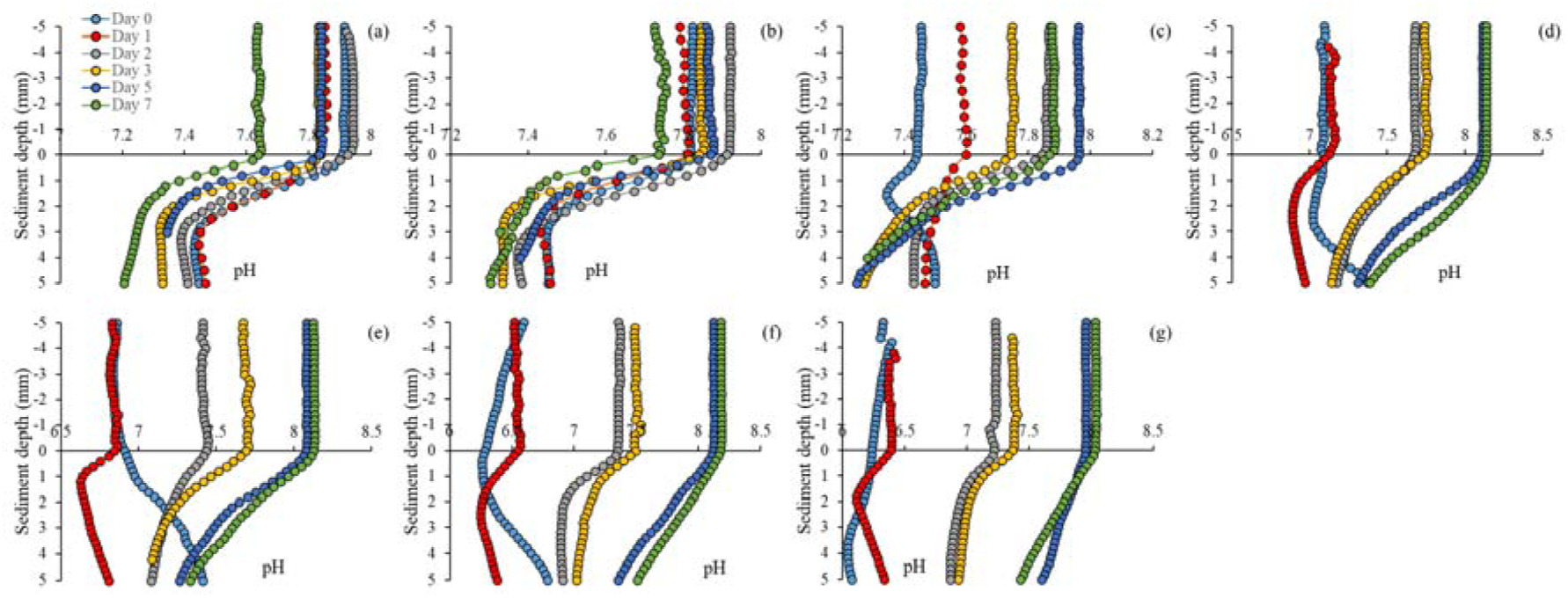
The pH depth profiles for the (a) DOC0, (b) DOC15, (c) DOC30, and (d) DOC45, (e) DOC60, (f) DOC75, and (g) DOC90 (day of culture). Values represent averages of biological triplicates, and error bars are omitted to maintain the visibility of the graphs. Zero depth equals to the sediment-water interface.

### 3.2. Effect of molybdate treatment in function of the shrimp culturing stage

Since there was still dissolved oxygen present at DOC30 and lower waste loads, indicating suitable conditions for shrimp growth, four different experiments were conducted to determine the effect of two different molybdate concentrations (25 and 50 mg L^-1^ Na_2_MoO_4_.2H_2_O) at different days of culture (DOC 45, 60 and 90) to control H_2_S formation. In all four treatments, the addition of molybdate, at both concentrations, at least partially suppressed H_2_S formation (Figure 4), but it did not impact the dissolved oxygen concentration (Figure 5) in the water column.

**Figure 4.**
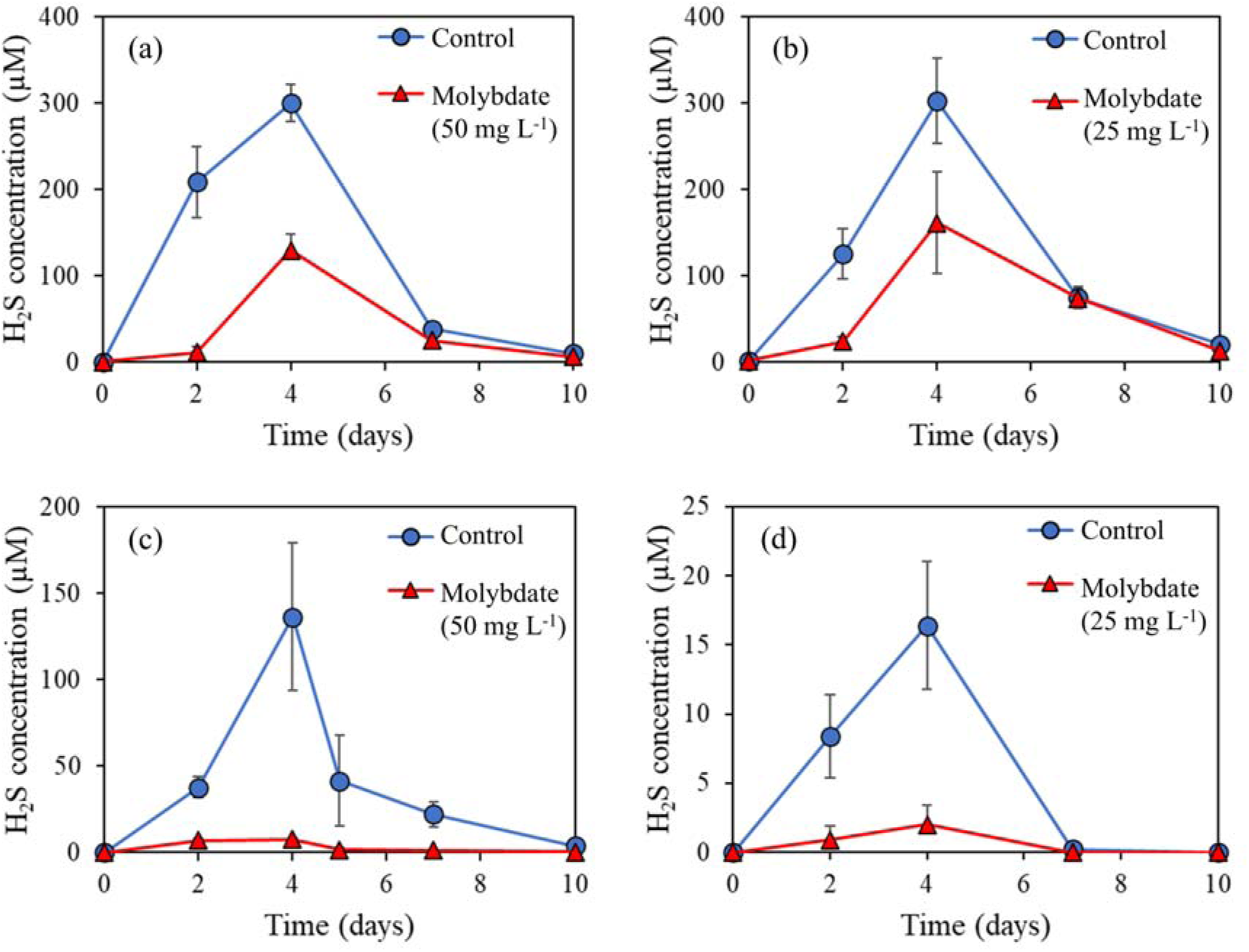
Water column H_2_S concentrations in the control and molybdate treatments for (a) DOC90 with 50 mg L^-1^ Na_2_MoO_4_.2H_2_O, (b) DOC90 with 25 mg L^-1^ Na_2_MoO_4_.2H_2_O, (c) DOC60 with 50 mg L^-1^ Na_2_MoO_4_.2H_2_O, and (d) DOC45 with 25 mg L^-1^ Na_2_MoO_4_.2H_2_O. Values represent averages of biological triplicates, and error bars represent standard deviations.

**Figure 5.**
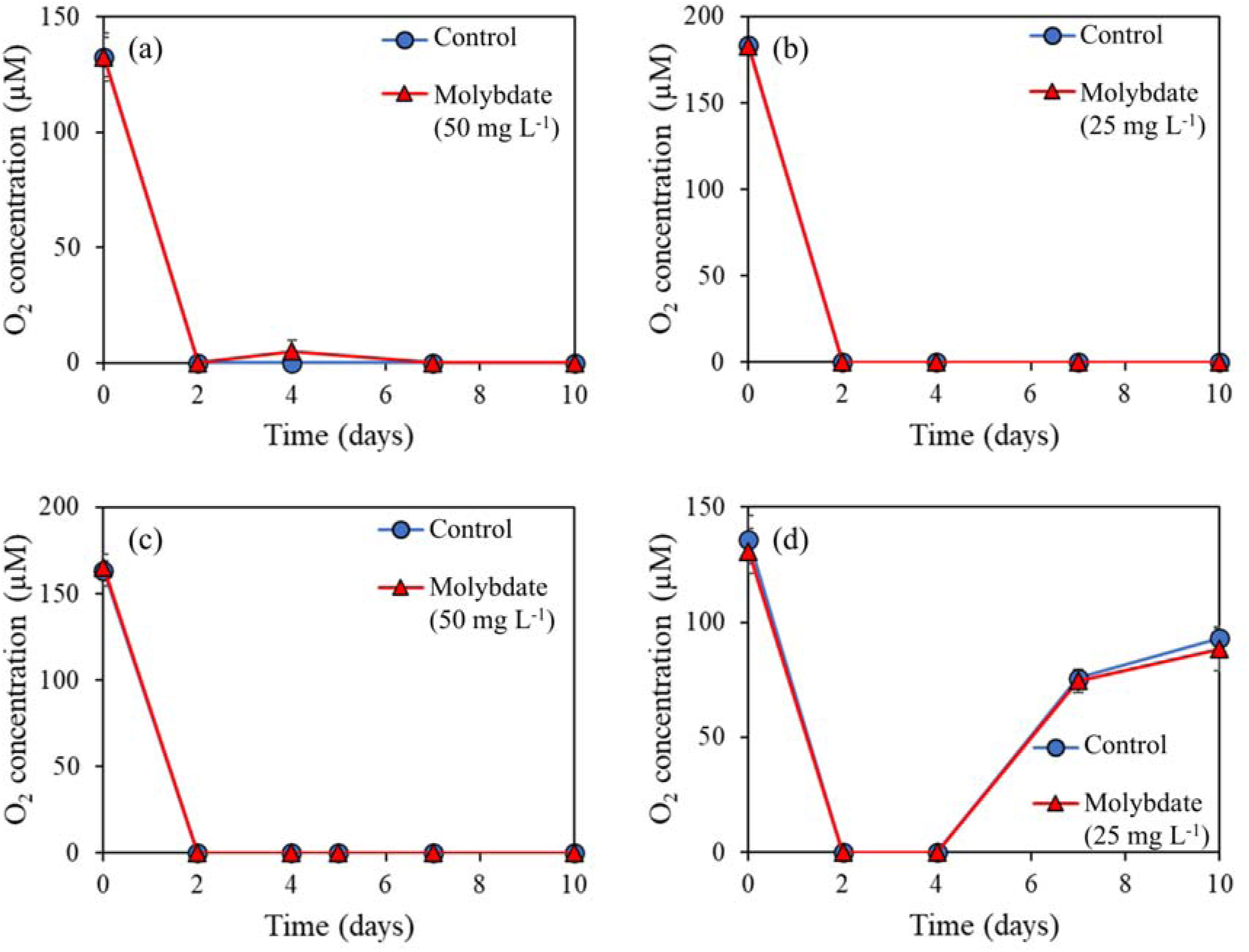
Water column O_2_ concentrations in the control and molybdate treatments for (a) DOC90 with 50 mg L^-1^ Na_2_MoO_4_.2H_2_O, (b) DOC90 with 25 mg L^-1^ Na_2_MoO_4_.2H_2_O, (c) DOC60 with 50 mg L^-1^ Na_2_MoO_4_.2H_2_O, and (d) DOC45 with 25 mg L^-1^ Na_2_MoO_4_.2H_2_O. Values represent averages of biological triplicates, and error bars represent standard deviations.

In the DOC90 treatment with 50 mg L^-1^ Na_2_MoO_4_.2H_2_O, there was a strong suppression of H_2_S formation on day 2, with a 95 ± 58 % lower H_2_S concentration in the water column compared to the control treatment (no molybdate added), and on day 4, with a 57 ± 9 % lower H_2_S concentration. However, while on day 2 the H_2_S formation was largely suppressed with an average value of 11 ± 6 µM, the H_2_S concentration increased to 129 ± 19 µM on day 4. On day 7 and day 10, despite a decline in H_2_S concentration in the water column, no clear difference between the control and molybdate treatment could be observed anymore. After day 0, oxygen could only be detected on day 4 in the molybdate treatment at a concentration of 4.9 ± 5.0 µM. A similar observation for H_2_S and dissolved oxygen could be made for the DOC90 treatment with 25 mg L^-1^ Na_2_MoO_4_.2H_2_O, yet, suppression of H_2_S formation on day 2 (81 ± 26 %) and day 4 (47 ± 19 %) was less pronounced compared to the 50 mg L^-1^ Na_2_MoO_4_.2H_2_O treatment.

Despite variations in the relative decrease of H_2_S, resulting from molybdate addition, the absolute reduction in H_2_S concentration was comparable between the molybdate-treated groups and the control in the DOC90 treatment. Specifically, in the DOC90 treatment with 50 mg L^-1^ Na_2_MoO_4_.2H_2_O, the absolute decrease in H_2_S was 171 ± 28 µM on day 4, while in the 25 mg L^-1^ Na_2_MoO_4_.2H_2_O treatment, it was 141 ± 77 µM.

The addition of molybdate appeared to have a stronger relative H_2_S suppression effect at lower waste loads in the DOC45 and DOC60 treatments. The addition of 50 mg L^-1^ Na_2_MoO_4_.2H_2_O to the DOC60 treatment resulted in a maximum H_2_S concentration of 7.7 ± 0.8 µM compared to 136 ± 43 µM in the control treatment on day 4, reflecting a 94 ± 31 % lower H_2_S concentration. A similar observation could be made towards the addition of 25 mg L^-1^ Na_2_MoO_4_.2H_2_O to the DOC45 treatment, which resulted in a maximum H_2_S concentration of 2.0 ± 1.4 µM, compared to 16.4 ± 4.6 µM in the control treatment on day 4, reflecting a 88 ± 67 % lower H_2_S production. Also here, for the DOC60 and DOC45, no difference could be observed in dissolved oxygen between the molybdate and control treatment, yet, in the DOC45 treatment, on day 7 and 10, oxygen was (passively) reintroduced in the system, which was not the case for the DOC60 treatment.

The pH showed a similar trend across various culture days and molybdate concentrations, including both the treatment and control groups (Figure S2). A consistent trend emerged, with an initial pH drop on day 2 in all treatments, followed by a subsequent increase. By day 10, the pH was higher than the initial pH observed on day 0.

The sulphate concentrations in the DOC90 treatment clearly decreased over time, both in the control and the molybdate treatment (Figure 6), however, the molybdate treatment had markedly higher residual sulphate concentrations than the control, both for 50 and 25 mg L^-1^ Na_2_MoO_4_.2H_2_O treatments. The molybdate treatment in DOC90 with 50 mg L^-1^ Na_2_MoO_4_.2H_2_O showed a 38 ± 1 % decrease in sulphate over time, and more sulphate was removed in the control treatment, reaching a 49 ± 4 % consumption of sulphate. Similar results were obtained in the DOC90 with 25 mg L^-1^ Na_2_MoO_4_.2H_2_O treatment. For the DOC60 treatment, there was a lower degree of sulphate reduction than the DOC90 treatment, and the molybdate treatment showed almost no decrease in sulphate over time (3.8 ± 0.0 %), compared to the control treatment (22 ± 1 %). In the DOC45 treatment, no substantial sulphate reduction was observed for both the control and molybdate treatment.

**Figure 6.**
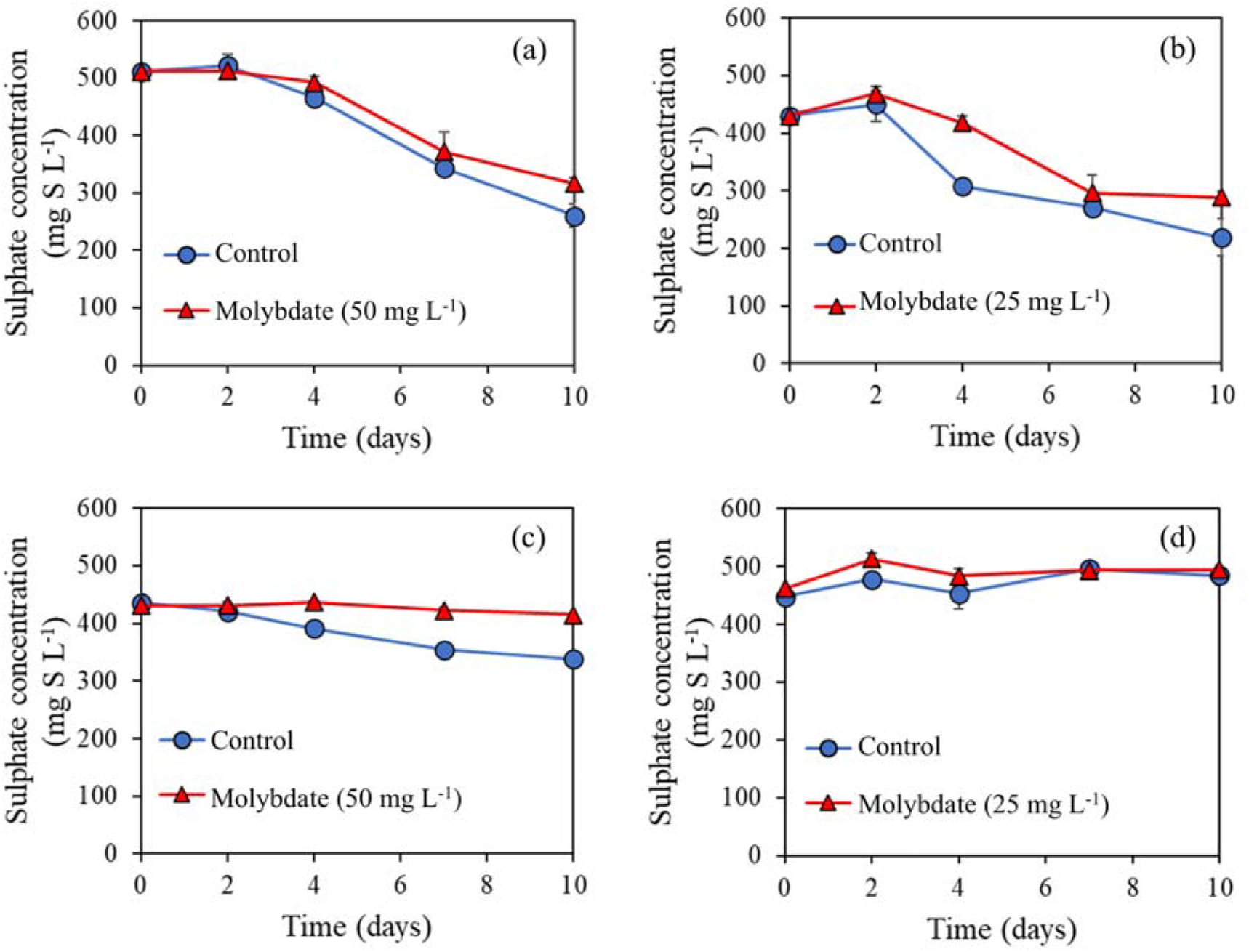
Water column sulphate concentrations in the control and molybdate treatments for (a) DOC90 with 50 mg L^-1^ Na_2_MoO_4_.2H_2_O, (b) DOC90 with 25 mg L^-1^ Na_2_MoO_4_.2H_2_O, (c) DOC60 with 50 mg L^-1^ Na_2_MoO_4_.2H_2_O, and (d) DOC45 with 25 mg L^-1^ Na_2_MoO_4_.2H_2_O. Values represent averages of biological triplicates, and error bars represent standard deviations.

The residual molybdate measurements showed that molybdate was fully reduced or utilized in both the DOC90 treatments at 50 and 25 mg L^-1^ Na_2_MoO_4_.2H_2_O (Table 1). At the lowest molybdate concentration, no molybdate was detected anymore on day 4, while this lasted till day 7 for the highest molybdate concentration. In the DOC60 and DOC45 treatments, no complete molybdate consumption occurred, and 11.6 ± 0.7 % and 66.4 ± 4.5 %, respectively, of molybdate remained unused at the end of the experiment. No molybdate was observed in any of the control treatments.

**Table 1.**
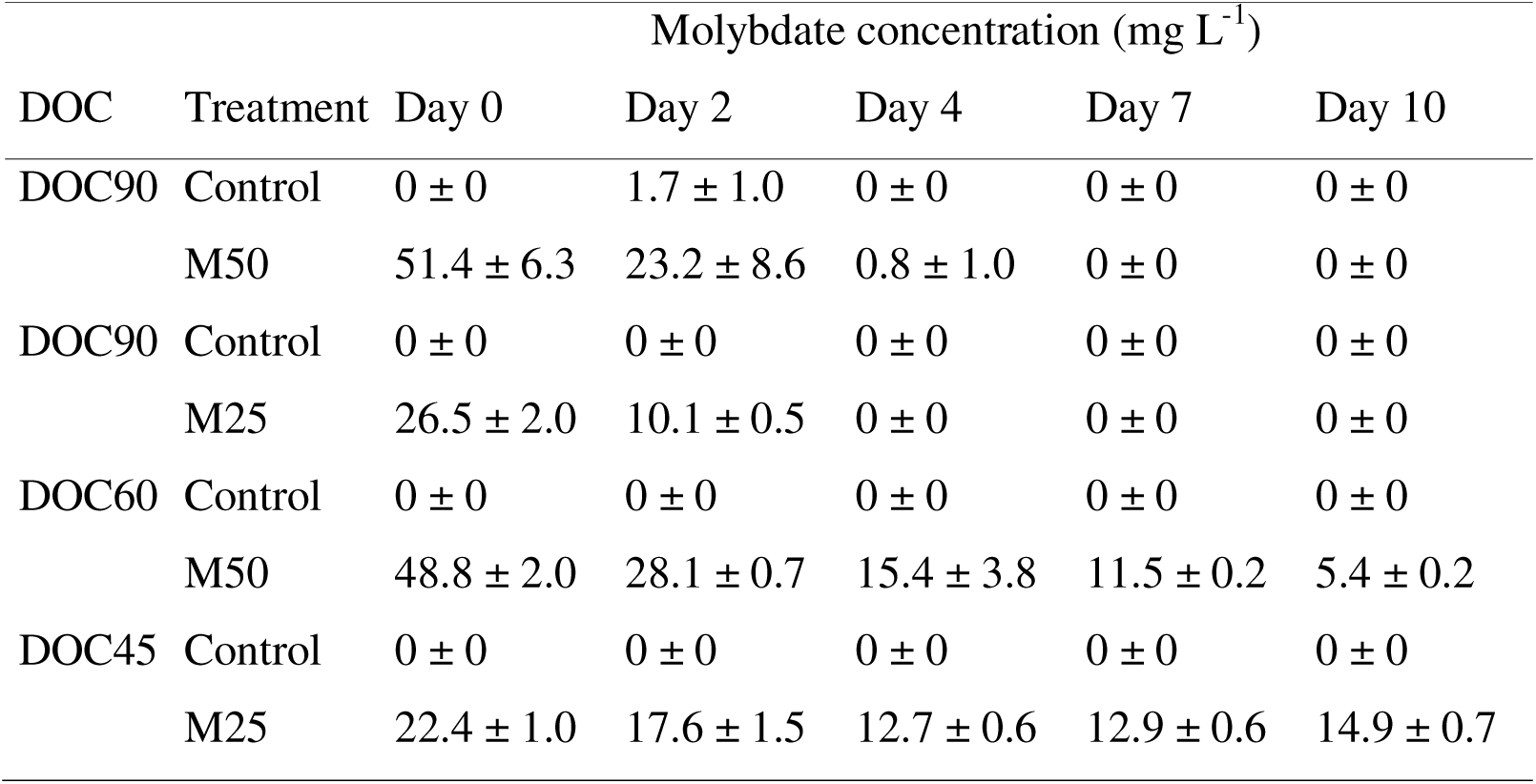
Molybdate concentrations of bulk liquid samples from the control and molybdate treatments at 25 (M25) and 50 mg L-1 Na2MoO4.2H2O (M50) at DOC90, DOC60 and DOC45 (day of culture). Values represent averages of biological triplicates with standard deviations.

## 4. Discussion

In this study, we evaluated the hypothesis that molybdate can be used as a preventive, targeted approach to suppress sulphide formation in shrimp pond bottoms at the appropriate moment in shrimp culturing. Oxygen depletion and sulphide accumulation took place in the middle of the 90-days shrimp growth period, *i.e.*, approximately around DOC45. The application of molybdate during the onset of sulphide production, *i.e.*, DOC45 and DOC60, resulted in a strong suppression of H_2_S formation. At heavy accumulation of organic waste (DOC90), molybdate still enabled the suppression of H_2_S formation, though less effective than at lower waste loads. No differences in dissolved oxygen were observed between the control and treatments, and only at DOC45, there was passive reintroduction of oxygen.

### 4.1. Oxygen depletion and sulphide production take place in the middle of the shrimp growth period

During the first 30 days of the 90-days shrimp growth period, oxygen depletion was limited, resulting in typical dissolved oxygen concentrations that remained in the range of 2.4-6.4 mg O_2_ L^-1^ (75-200 µM O_2_), hence, well within or at least close to the minimum oxygen concentration range (3-4 mg O_2_ L^-1^) that is recommended for shrimp dwelling on the pond bottom (Boyd and Hanson, 2010). Since the intrusion of oxygen in the laminar boundary layer close to the sediment and in the sediment itself is limited to transportation via diffusion, the (re)aeration of the sediment layer is a slow process. Hence, once the consumption of oxygen exceeds the rate of oxygen supply, oxygen is depleted, and this is exactly what was observed starting from DOC45 on in our experiments. In this study, oxygen diffusion was measured in the first 4 mm of the upper sediment layer in the DOC0 and DOC15 treatment, yet, in the DOC30 treatment, this penetration depth decreased to 3 mm, despite leaving sufficient oxygen in the water column where the shrimp dwell. On DOC45, this penetration depth further decreased, severely impacting the dissolved oxygen in the water column, resulting in anaerobic conditions. These results are in line with research where oxygen penetration depth in semi-intensive fish pond sediments was limited to 1 mm (Meijer and Avnimelech, 1999). The depletion of oxygen in the water column just above the sediment has a double negative impact towards the shrimp dwelling there. First, shrimp can survive anaerobic conditions for only a few minutes (Boyd and Hanson, 2010), and our results have shown that already from DOC45 on, oxygen is depleted for at least several days in a row. Second, the presence of sulphate and the activity of sulphate reducing bacteria provoke the formation of toxic H_2_S, which can be released from the sediment into the water column through diffusion (Avnimelech and Ritvo, 2003). Like for oxygen, but in the opposite direction, this diffusion-limited transport of H_2_S from the sediment towards the water column is delayed. This was observed in our experiments, amongst others in DOC45, where on day 2 of this treatment the H_2_S concentration remained below 1 µM in the water column, yet, reaching up to 12 µM in the sediment. A subsequent increase in H_2_S in the water column was only observed in day 3 of DOC45. This observation confirms that H_2_S formation mainly takes place in the sediment and not the water column, and was most likely due the selective enrichment of sulphate reducing bacteria. As observed in our results, the rapid depletion of oxygen in the sediment in function of depth creates the ideal conditions for the obligatory anaerobic sulphate reducing bacteria, in contrast to the water column where oxygen lingers and/or reaeration can take place, as observed on day 5 and 7 in DOC45.

About 50-80% of total organic carbon degradation in shrimp pond sediments was reported to take place through non-aerobic processes, such as nitrate, iron, and manganese reduction, but also sulphate reduction and methanogenesis (Burford et al., 1998). This coincides with our research findings that sulphate reduction in shrimp pond sediments is a key process towards the biodegradation of organic matter. Hence, there is a substantial amount of organic matter, taking into consideration the intensive nature of shrimp growth, that requires dealing with to avoid both oxygen limitation and H_2_S intrusion in the water column.

### 4.2. Molybdate can act as a suitable strategy to suppress H_2_S formation when applied at an early stage in the shrimp growth period

As mentioned earlier, an effective strategy that prevents both mortality and sub-lethal effects on shrimp needs to ensure that oxygen is not depleted, should avoid or at least limit the intrusion of H_2_S in the water column, but should also avoid the introduction of novel toxicants, which is in line with avoiding the “law of conservation of misery”. In this study, molybdate was chosen as a potential strategy to suppress H_2_S formation. Concentrations of 25 and 50 mg L^-1^ molybdate were used, as these were previously shown to be effective in reducing H_2_S formation in shrimp pond bottom models (Torun et al., 2022; Torun et al., 2024). We considered the DOC treatments with minor (DOC45 and DOC60) and severe (DOC90) organic matter accumulation and H_2_S formation during shrimp growth, as determined in this study. This was done to evaluate the potential of molybdate as an effective H_2_S suppression strategy.

In the DOC45 and DOC60 treatments, H_2_S formation was effectively suppressed, based on both the H_2_S concentration and residual sulphate concentration in the liquid. In contrast, both DOC90 treatments also showed a clear decrease in H_2_S formation, yet, after day 0, a rapid increase was again observed to values in the order of 5 mg L^-1^ (150 µM), which is well above the levels of 0.02 mg L^-1^ causing already toxicity to shrimp (US-EPA, 2011). In the DOC45 and DOC60 treatments, there was also still residual molybdate left, which was not the case in the DOC90 treatments. In addition, since in all treatments there was still residual sulphate left, it appears that the presence of biodegradable organics is the key limiting factor towards sulphate reduction and H_2_S formation, and molybdate is only effective in earlier stages of the shrimp growth cycle. The almost complete consumption of molybdate on day 4 in the DOC90 treatments aligns with a spike in H_2_S in the water column, indicating that the suppressive effect of molybdate was only temporal. This is in line with another study, where it was shown that molybdate was only bacteriostatic to sulphate reducing bacteria (Isa and Anderson, 2005). It appears that there is a wide variety of molybdate (MoO_4_) to sulphate (SO_4_) ratios that are minimally needed to inhibit sulphate reduction, with complete inhibition observed at a molybdate to sulphate ratio of 1:3 (Chen et al., 1998). Partial inhibition was observed in one study at a molybdate to sulphate ratio of 1:190 (Chen et al., 1998), while in another study a molybdate to sulphate ratio of 1:250 was sufficient to suppress sulphate reduction for at least 168 hours (Jesus et al., 2015). In our study, the treatments with 50 and 25 mg L^-1^ Na_2_MoO_4_.2H_2_O roughly corresponded with a molybdate to sulphate ratio of 90 and 45 respectively, but the organic matter content, determined by the day of culture, rather than the molybdate to sulphate ratio, determined the effectiveness of the molybdate treatment. Hence, the sustained presence of residual molybdate, rather than its actual concentration or the molybdate to sulphate ratio, is essential to sustaining the suppression of sulphate reduction, and thus H_2_S formation.

The presence of residual molybdate might create another problem, as mentioned earlier, reflecting the law of conservation of misery. A comprehensive study by De Schamphelaere et al. (2010), studying the EC_10_ (concentration effecting 10% of species for their most sensitive endpoint) of molybdate for 10 different freshwater aquatic species (both vertebrates and invertebrates), showed a high variability between species, yet, a median hazardous concentration affecting 5% of the species of 38.2 mg L^-1^ molybdenum, corresponding to 96 mg L^-1^ Na_2_MoO_4_.2H_2_O, which is about twice as high as the maximal concentration applied in our study. A similar study in which nine marine species, representative for typical marine trophic levels, were chronically exposed to molybdate also showed a high degree of variation, yet, in this case a median hazardous concentration affecting 5% of the species of 5.74 mg L^-1^ molybdenum, corresponding to 14.5 mg L^-1^ Na_2_MoO_4_.2H_2_O, was observed (Heijerick et al., 2012). However, in this same study, for the *Americamysis bahia*, another shrimp-like crustacean, an EC_10_ value > 116 mg L^-1^ molybdenum was observed (Heijerick et al., 2012). Hence, it appears that the presence of residual molybdate at the concentrations measured in our research is not directly toxic to shrimp, yet specific long-term toxicity tests with *Litopenaeus vannamei* should be carried out to confirm this. Certain marine species, like the blue mussel (*Mytilus edulis*), appear to be sensitive, with an EC_10_ value of 4.40 mg L^-1^ molybdenum (Heijerick et al., 2012), corresponding to 11.1 mg L^-1^ Na_2_MoO_4_.2H_2_O, which falls within the range of the concentrations applied and measured in our study.

The application of molybdate, especially at early phases in the shrimp growth cycle could successfully suppress H_2_S formation, yet, it did not promote the (re)introduction of oxygen, as no difference in dissolved O_2_ concentration in the water column were observed in any of the four molybdate treatments between the control and molybdate treatment. Molybdate plays a key role in both sulphur and nitrogen cycling in almost all living organisms (Huang et al., 2021), yet, it does not directly appear to play a key role in oxygen uptake. Hence, based on our results, it implies that the application of molybdate as such is insufficient to ensure the reintroduction of oxygen, and other, more active forms of aeration, such as mechanical aeration are also needed to ensure steady oxygen levels in the water column.

## 5. Conclusions

We showed that both oxygen depletion and sulphide formation take place in a shrimp pond bottom model in the water column in the middle of the shrimp growth period. Molybdate addition was effective in suppressing sulphide formation only in the early stages of the shrimp growth period. The presence of residual molybdate appeared to be essential to sustain sulphate reduction suppression, irrespective of the sulphate concentration, yet, potential toxicity to marine life warrants caution. While molybdate effectively controlled sulphide formation, it did not result in an increased dissolved oxygen concentration, implying the need for additional aeration strategies to sustain oxygen levels.

## Acknowledgments

Special thanks go to the Laboratory of Aquaculture and Artemia Reference Center, Ghent University, for providing shrimp feed (Crevetec Grower 2) and faeces. We would like to thank Koenraad Maréchal and Dirk Raes from Agentschap Natuur & Bos (Belgium) for their help during collection of sediment samples.

## Data, script, code and supplementary material

The datasets generated during this research are included in this article, its supplementary information, and were submitted to the Zenodo repository (https://zenodo.org/doi/10.5281/zenodo.13359841).

## References

Alonso-Rodrı guez, R. and Páez-Osuna, F. 2003. Nutrients, phytoplankton and harmful algal blooms in shrimp ponds: a review with special reference to the situation in the Gulf of California. Aquaculture 219(1), 317–336.

Avnimelech, Y. and Ritvo, G. 2003. Shrimp and fish pond soils: processes and management. Aquaculture 220(1), 549–567.

Boyd, C. and Hanson, T. 2010. Dissolved-oxygen concentration in pond aquaculture. Global Aquacult. Advocate 13, 40–41.

Boyd, C.E. 1998. Pond water aeration systems. Aquacultural Engineering 18(1), 9–40.

Burford, M.A., Peterson, E.L., Baiano, J.C.F. and Preston, N.P. 1998. Bacteria in shrimp pond sediments: their role in mineralizing nutrients and some suggested sampling strategies. Aquaculture Research 29(11), 843–849.

Chen, G., Ford, T.E. and Clayton, C.R. 1998. Interaction of Sulfate-Reducing Bacteria with Molybdenum Dissolved from Sputter-Deposited Molybdenum Thin Films and Pure Molybdenum Powder. J. Colloid Interface Sci. 204(2), 237–246.

Ciji, A. and Akhtar, M.S. 2020. Nitrite implications and its management strategies in aquaculture: a review. Reviews in Aquaculture 12(2), 878–908.

De Schamphelaere, K.A.C., Stubblefield, W., Rodriguez, P., Vleminckx, K. and Janssen, C.R. 2010. The chronic toxicity of molybdate to freshwater organisms. I. Generating reliable effects data. Science of The Total Environment 408(22), 5362–5371.

Dien, L.D., Hiep, L.H., Faggotter, S.J., Chen, C., Sammut, J. and Burford, M.A. 2019. Factors driving low oxygen conditions in integrated rice-shrimp ponds. Aquaculture 512, 734315.

Greenberg, A.E., Clesceri, L.S. and Eaton, A.D. 1992. Standard Methods for the Examination of Water and Wastewater American Public Health Association Publications, Washington.

Heijerick, D.G., Regoli, L. and Stubblefield, W. 2012. The chronic toxicity of molybdate to marine organisms. I. Generating reliable effects data. Science of The Total Environment 430, 260–269.

Huang, X.-Y., Hu, D.-W. and Zhao, F.-J. 2021. Molybdenum: More than an essential element. Journal of Experimental Botany 73(6), 1766–1774.

Hung, L.t. and Quy, O.M. 2014. On-farm feeding and feed management in whiteleg shrimp (Litopenaeus vannamei) farming in Viet Nam.

Isa, M.H. and Anderson, G.K. 2005. Molybdate inhibition of sulphate reduction in two-phase anaerobic digestion. Process Biochemistry 40(6), 2079–2089.

Jesus, E., Lima, L., Bernardez, L.A. and Almeida, P. 2015. Inhibition of microbial sulfate reduction by molybdate. Brazilian Journal of Petroleum & Gas 9, 95.

Kannan, B., Felix, N., Rathipriya, A., Meeran, M.N. and Prabu, E. 2018. A review on pond botttom management in shrimp aquaculture. Journal of Aquaculture in the Tropics 33(3/4), 135–142.

Malkin, S.Y., Rao, A.M.F., Seitaj, D., Vasquez-Cardenas, D., Zetsche, E.-M., Hidalgo-Martinez, S., Boschker, H.T.S. and Meysman, F.J.R. 2014. Natural occurrence of microbial sulphur oxidation by long-range electron transport in the seafloor. Isme j 8(9), 1843–1854.

Meijer, L.E. and Avnimelech, Y. 1999. On the use of micro-electrodes in fish pond sediments. Aquacultural Engineering 21(2), 71–83.

Nemati, M., Mazutinec, T.J., Jenneman, G.E. and Voordouw, G. 2001. Control of biogenic H_2_S production with nitrite and molybdate. Journal of Industrial Microbiology and Biotechnology 26(6), 350–355.

Panakorn, S. 2016. Hydrogen sulfide-The silent killer. Aquaculture Asia Pacific Magazine (March-April), 14–17.

Predicala, B., Nemati, M., Stade, S. and Laguë, C. 2008. Control of H_2_S emission from swine manure using Na-nitrite and Na-molybdate. Journal of Hazardous Materials 154(1), 300–309.

Schwermer, C.U., Ferdelman, T.G., Stief, P., Gieseke, A., Rezakhani, N., Van Rijn, J., De Beer, D. and Schramm, A. 2010. Effect of nitrate on sulfur transformations in sulfidogenic sludge of a marine aquaculture biofilter. FEMS Microbiology Ecology 72(3), 476–484.

Shiau, S.-Y. 1998. Nutrient requirements of penaeid shrimps. Aquaculture 164(1), 77–93.

Suo, Y., Li, E., Li, T., Jia, Y., Qin, J.G., Gu, Z. and Chen, L. 2017. Response of gut health and microbiota to sulfide exposure in Pacific white shrimp *Litopenaeus vannamei*. Fish Shellfish Immunol 63, 87–96.

Tang, K., Baskaran, V. and Nemati, M. 2009. Bacteria of the sulphur cycle: An overview of microbiology, biokinetics and their role in petroleum and mining industries. Biochemical Engineering Journal 44(1), 73–94.

Thunjai, T., Boyd, C.E. and Dube, K. 2001. Poind Soil pH Measurement. Journal of the World Aquaculture Society 32(2), 141–152.

Torun, F., Hostins, B., De Schryver, P., Boon, N. and De Vrieze, J. 2022. Molybdate effectively controls sulphide production in a shrimp pond model. Environmental Research 203, 111797.

Torun, F., Hostins, B., De Schryver, P., Boon, N. and De Vrieze, J. 2024. Molybdate delays sulphide formation in the sediment and transfer to the bulk liquid in a model shrimp pond. Peer Community Journal 4.

Torun, F., Hostins, B., Teske, J., De Schryver, P., Boon, N. and De Vrieze, J. 2020. Nitrate amendment to control sulphide accumulation in shrimp ponds. Aquaculture 521, 735010.

US-EPA 2011. Hydrogen Sulfide; Community Right-to-Know Toxic Chemical Release Reporting, pp. 64022–64037.

Vismann, B. 1996. Sulfide species and total sulfide toxicity in the shrimp *Crangon crangon*. Journal of Experimental Marine Biology and Ecology 204(1), 141–154.

Will, F. and Yoe, J.H. 1953. Colorimetric Determination of Molybdenum with Mercaptoacetic Acid. Analytical Chemistry 25, 1363–1366.

