## Supplementary file 1 for "Molybdate application in the early stages of shrimp growth suppresses sulphide formation in a shrimp pond bottom model"

Journal: bioRxiv

### S1. Experimental set-up

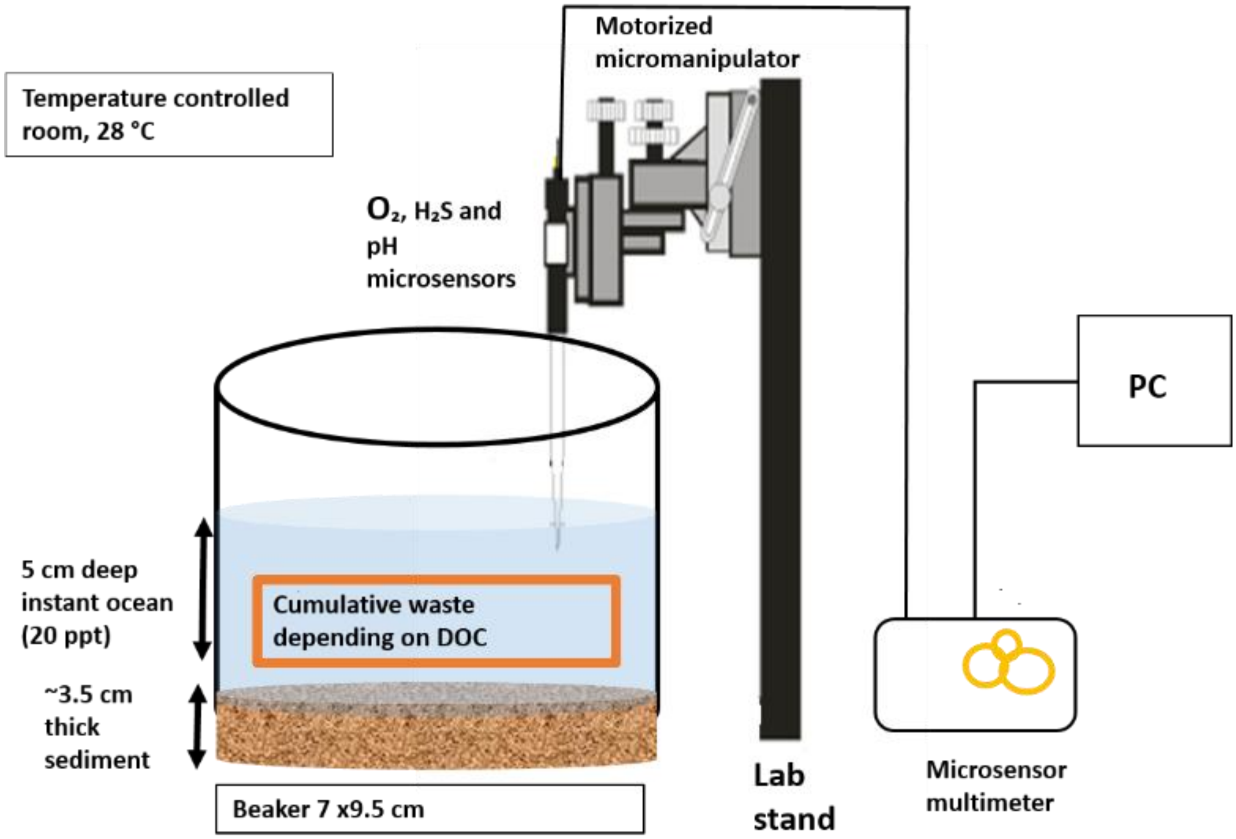

**Figure S1.** The experimental shrimp pond model, including the sediment layer and overlaying artificial seawater. Sediment microscale depth profiles of $O_2$, $H_2S$ and pH were determined by the use of microelectrodes and a microprofiling set-up (Unisense, Denmark). The microprofiling set-up has a motorized micromanipulator that helps to move the microelectrodes at micrometer-size step through the sediment depth and the microsensor multimeter that acts as amplifier at which the microelectrodes could be plugged in at their correct polarization value.

#### S2. Shrimp feeding tables

**Table S1** Daily feeding table for the shrimp post-larvae (PL) stocked at 50 shrimp per $m^3$ density

| **Day of culture (DOC)** | **PL stage** | **PL size (mg)** | **Daily feeding rate (% of PL mass)** | **Required feed ($g\ m^{-3}\ d^{-1}$)** |
|---|---|---|---|---|
| 0 | 10 | 6 | 34 | 0.102 |
| 1 | 11 | 7 | 33 | 0.107 |
| 2 | 12 | 7 | 32 | 0.112 |
| 3 | 13 | 11 | 31 | 0.171 |
| 4 | 14 | 15 | 30 | 0.225 |
| 5 | 15 | 20 | 29 | 0.29 |
| 6 | 16 | 25 | 28 | 0.35 |
| 7 | 17 | 29 | 28 | 0.406 |
| 8 | 18 | 33 | 27 | 0.446 |
| 9 | 19 | 38 | 27 | 0.513 |
| 10 | 20 | 45 | 27 | 0.608 |
| 11 | 21 | 60 | 26 | 0.780 |
| 12 | 22 | 80 | 26 | 1.040 |
| 13 | 23 | 100 | 26 | 1.300 |
| 14 | 24 | 105 | 25 | 1.312 |
| 15 | 25 | 110 | 25 | 1.375 |
| 16 | 26 | 135 | 22 | 1.485 |
| 17 | 27 | 155 | 20 | 1.550 |
| 18 | 28 | 185 | 20 | 1.850 |
| 19 | 29 | 215 | 19 | 2.043 |
| 20 | 30 | 245 | 19 | 2.328 |
| 21 | 31 | 275 | 18 | 2.475 |
| 22 | 32 | 310 | 18 | 2.790 |
| 23 | 33 | 345 | 18 | 3.105 |
| 24 | 34 | 400 | 17 | 3.400 |
| 25 | 35 | 465 | 17 | 3.953 |
| 26 | 36 | 550 | 17 | 4.675 |
| 27 | 37 | 640 | 16 | 5.120 |
| 28 | 38 | 695 | 16 | 5.560 |
| 29 | 39 | 750 | 16 | 6.000 |
| 30 | 40 | 835 | 15 | 6.263 |
| 31 | 41 | 920 | 15 | 6.900 |
| 32 | 42 | 1025 | 15 | 7.688 |
| 33 | 43 | 1130 | 14 | 7.910 |
| 34 | 44 | 1240 | 14 | 8.680 |
| 35 | 45 | 1360 | 14 | 9.520 |
| 36 | 46 | 1480 | 14 | 1.360 |
| 37 | 47 | 1600 | 13 | 10.400 |

| | | | | |
|---|---|---|---|---|
| 38 | 48 | 1720 | 13 | 11.180 |
| 39 | 49 | 1850 | 13 | 12.025 |
| 40 | 50 | 1970 | 13 | 12.805 |
| 41 | 51 | 2100 | 12 | 12.600 |
| 42 | 52 | 2250 | 12 | 13.500 |
| 43 | 53 | 2340 | 12 | 14.040 |
| 44 | 54 | 2434 | 12 | 14.602 |
| 45 | 55 | 2531 | 11 | 13.920 |
| 46 | 56 | 2632 | 11 | 14.477 |
| 47 | 57 | 2737 | 11 | 15.056 |
| 48 | 58 | 2847 | 11 | 15.658 |
| 49 | 59 | 2961 | 10 | 14.804 |
| 50 | 60 | 3079 | 10 | 15.396 |
| 51 | 61 | 3202 | 10 | 16.012 |
| 52 | 62 | 3331 | 10 | 16.658 |
| 53 | 63 | 3464 | 10 | 17.319 |
| 54 | 64 | 3602 | 9 | 16.211 |
| 55 | 65 | 3746 | 9 | 16.859 |
| 56 | 66 | 3896 | 9 | 17.534 |
| 57 | 67 | 4052 | 9 | 18.235 |
| 58 | 68 | 4214 | 9 | 18.964 |
| 59 | 69 | 4383 | 8 | 17.531 |
| 60 | 70 | 4558 | 8 | 18.232 |
| 61 | 71 | 4740 | 8 | 18.961 |
| 62 | 72 | 4930 | 8 | 19.720 |
| 63 | 73 | 5127 | 8 | 20.509 |
| 64 | 74 | 5332 | 7 | 18.663 |
| 65 | 75 | 5546 | 7 | 19.410 |
| 66 | 76 | 5767 | 7 | 20.186 |
| 67 | 77 | 5998 | 7 | 20.993 |
| 68 | 78 | 6238 | 7 | 21.833 |
| 69 | 79 | 6488 | 6 | 19.463 |
| 70 | 80 | 6747 | 6 | 20.241 |
| 71 | 81 | 7017 | 6 | 21.051 |
| 72 | 82 | 7298 | 6 | 21.893 |
| 73 | 83 | 7590 | 6 | 22.769 |
| 74 | 84 | 7893 | 5 | 19.733 |
| 75 | 85 | 8209 | 5 | 20.522 |
| 76 | 86 | 8537 | 5 | 21.343 |
| 77 | 87 | 8879 | 5 | 22.197 |
| 78 | 88 | 9234 | 5 | 23.085 |
| 79 | 89 | 9603 | 4 | 19.206 |
| 80 | 90 | 9987 | 4 | 19.975 |
| 81 | 91 | 10387 | 4 | 20.774 |
| 82 | 92 | 10802 | 4 | 21.605 |

| | | | | |
|---|---|---|---|---|
| 83 | 93 | 11234 | 4 | 22.469 |
| 84 | 94 | 11684 | 4.5 | 26.288 |
| 85 | 95 | 12151 | 4.5 | 27.340 |
| 86 | 96 | 12637 | 4.5 | 28.433 |
| 87 | 97 | 13143 | 4.5 | 29.570 |
| 88 | 98 | 13668 | 4.5 | 30.754 |
| 89 | 99 | 14215 | 3.5 | 24.876 |
| 90 | 100 | 14784 | 3.5 | 25.871 |

#### S3. Organic waste supplementation table

**Table S2** The amount of organic waste (feed and faeces) supplemented in the different treatments. The amount of feed and faeces were determined to account for the increasing uneaten feed and faeces accumulation on the pond bottom with the increasing size of shrimp, due to growth. The volume of faeces supplemented was determined based on its dry matter content (4.4%), while the amount of feed added was determined based on Table S1.

| **Day of culture** | **Feed supplement (mg day$^{-1}$)** | **Faeces supplement (mL day$^{-1}$)** |
|---|---|---|
| DOC0 | 0 | 0 |
| DOC15 | 4 | 0.12 |
| DOC30 | 27 | 0.81 |
| DOC45 | 99.7 | 2.99 |
| DOC60 | 208.5 | 6.26 |
| DOC75 | 342.3 | 10.27 |
| DOC90 | 501.4 | 15.04 |

#### S4. Water column pH

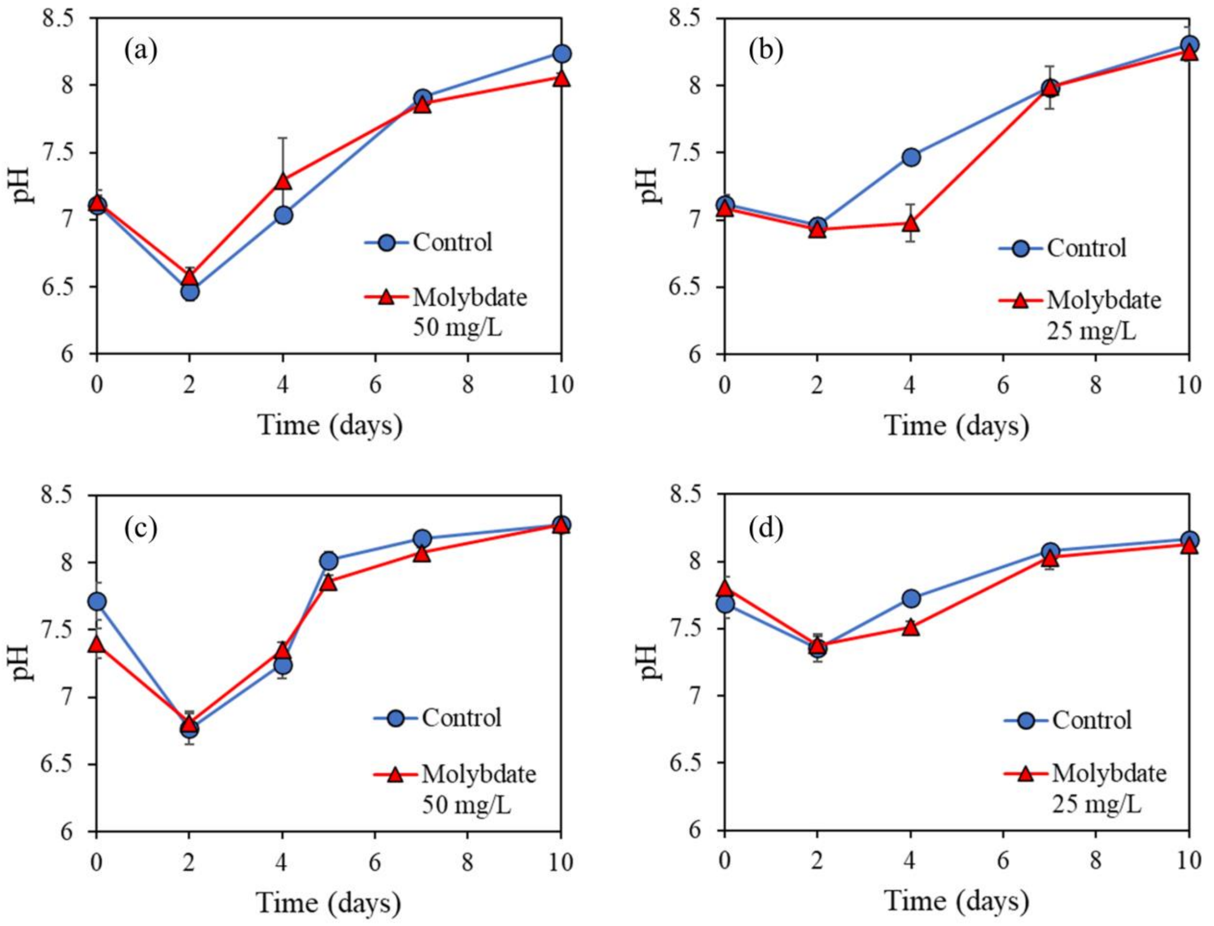

**Figure S2** Water column pH values in the control and molybdate treatments for (q) DOC90 with 50 mg $L^{-1}$ $Na_2MoO_4.2H_2O$, (b) DOC90 with 25 mg $L^{-1}$ $Na_2MoO_4.2H_2O$, (c) DOC60 with 50 mg $L^{-1}$ $Na_2MoO_4.2H_2O$, and (d) DOC45 with 25 mg $L^{-1}$ $Na_2MoO_4.2H_2O$. Values represent averages of biological triplicates, and error bars represent standard deviations.
